# An updated framework to account for inter-individual variability when quantifying phenotypic variation

**DOI:** 10.1101/2021.10.22.465436

**Authors:** Giacomo Puglielli, Carlos P. Carmona, Laura Varone, Lauri Laanisto, Carlo Ricotta

## Abstract

1. In trait-based ecology, phenotypic variation (*PVar*) is often quantified with measures that express average differences between populations standardized in the range 0-1. A major problem with these measures is that they disregard the within-population trait variability. In addition, most of these measures cannot be decomposed across scales. This can alter their interpretation, thus limiting their applicability.
2. To overcome these problems, we propose a new measure, the Phenotypic Dissimilarity Index (PhD) that is insensitive to the within-population interindividual trait variability. Likewise, PhD can be used to quantify *PVar* between individuals in a population while accounting for the *PVar* within individuals.
3. Using simulated and real data, we showed that PhD index correctly quantifies *PVar* when the within-population trait variability is not negligible, as in many ecological studies. By accounting for within-population trait variability, the PhD index generally provides a more parsimonious quantification of *PVar* across an environmental gradient compared to other estimators.
4. Traits sampled within a species have an inherent variability. Accounting for such variability is essential to understand species phenotypic responses to environmental cues. As such, the PhD index will provide ecologists with an asset to reliably quantify and compare *PVar* within and between species across environmental gradients at different scales. We also provide an R function to calculate the PhD index.

## Introduction

Functional traits variability within species is a key determinant of species’ ability to cope, and eventually adapt, to environmental factor variations. Accounting for this variability is considered essential to explain the success of a species under contrasting environmental conditions (Garnier et al., 2015). However, trait variability within species, or intraspecific trait variability (ITV), or phenotypic plasticity (*sensu* Valladares et al., 2000), or phenotypic variation, has been generally overlooked in the past as it was considered to be negligible compared to interspecific variability (Garnier et al., 2001; McGill et al., 2006). While this assumption holds true at the global scale, in the last years many studies have spotlighted the adaptive role of ITV for organismal responses to environmental changes at different scales (Arnold et al., 2019; Henn et al., 2018; Kuppler et al., 2020; Siefert et al., 2015; Wong & Carmona, 2021). As a result, there has been an increasing interest in developing methodological approaches to quantify ITV in functional trait studies (e.g. de Bello et al., 2011; Niu et al., 2020).

One of the major challenges in developing methods to quantify ITV in functional trait studies is the difficulty to include all its underlying sources and structure. ITV sources are genetic variability, phenotypic plasticity, and their interaction (Albert et al., 2011). However, differentiating between sources of ITV requires exact knowledge on individuals’ genotype, more than often lacking in most of the functional traits-based studies. On the other hand, ITV structure includes three levels of phenotypic variation among individuals of a species: (i) Differences between populations of a species – e.g. plastic responses of a given genotype to different static (Sandquist & Ehleringer, 1997) or temporally dynamic (e.g. Turner et al., 2008) environments; (ii) Inter-individual variability (Bolnick et al., 2003), defined as the trait variability among individuals within a population/sub-population – e.g. phenotypic variation of individual traits across micro-environmental conditions within a site/population; (iii) Within-individual variability (Bolnick et al., 2003; Herrera et al., 2015) – e.g. sun *vs*. shade leaves differences within a canopy (e.g. Niinemets et al., 2015), ontogenetically changing traits (Poorter et al., 2015; Puglielli et al., 2021), or seasonally variable traits (e.g. Mason et al., 2020; Puglielli, 2019; Puglielli et al., 2019).

Two methods are widely employed to quantify ITV: i) the reaction norm: a measure of the extent and direction of an environmentally induced phenotypic change (Arnold et al., 2019; Nicotra et al., 2010; Schlichting & Pigliucci, 1998); ii) various ‘indexes of phenotypic variation’ (*PVar*), reviewed by Valladares et al., 2006. We refer to such indexes as (*PVar*), instead of ‘phenotypic plasticity indexes’ as in Valladares et al. (2006), to clarify that they cannot discriminate between the sources of ITV, and thus they only quantify the phenotypic but not the genetic component of ITV (genetic information on the individuals under study are in fact rarely available). Here we focus on indexes of ‘phenotypic variation’ (*PVar*), and refer to the recent work of Arnold et al. (2019) for a recent review on possibilities and pitfalls of reaction norms (e.g. linearity *vs*. non-linearity of trait changes along gradients). A key advantage of using *PVar* when quantifying ITV is that they are relatively simple to calculate, and they provide estimates of phenotypic variation that can be easily compared among species and from which to draw straightforward ecological and evolutionary interpretations (e.g., Valladares et al., 2006). Among the many available *PVar*, the most used remain:

1. The Plasticity Index (PI) proposed by Valladares et al. (2000). PI is calculated as the difference between the maximum mean (M1) and minimum mean (M2) values of a trait in different environmental conditions divided by the maximum mean value (e.g. (M1 – M2)/M1; Balaguer et al., 2001; Castro-Díez et al., 2006; Puglielli et al., 2017; Rutherford et al., 2017). The major limitation of PI is that it quantifies only the between population trait variability component of ITV, totally disregarding the within population trait variability component (**Fig. 1a**). In doing that, mean differences between populations can be biased if some individuals within a population contribute to the population mean to a greater extent compared to other individuals.
2. The coefficient of variation (CV), calculated as the ratio between standard deviation and the mean of a given trait value in response to different environmental conditions. CV takes into account within population trait variability, as standard deviation is necessary for its calculation, but it cannot be decomposed across scales (**Fig. 1b**).
3. Relative Distance Plasticity Index (RDPI) (Valladares et al., 2006). This index calculates the mean phenotypic dissimilarity among all the individuals of a given species exposed to different environmental conditions. The advantage of using RDPI over the classic PI consists in the RDPI ability to express an average phenotypic distance among all the measured individuals between individuals of a given species growing in different environmental conditions (Valladares et al., 2006). While RPDI is an individual-based *PVar* that could in principle account for both between and within population trait variability component of ITV, here we show that RDPI index is limited because it does not account for intrapopulation variability (**Fig. 1c**), and because of this, RDPI values strongly depend on the trait variability within each group under comparison. In other words, by not accounting for within population trait variability when calculating between population trait variability, RDPI incurs into the same pitfall as PI.

**Fig. 1.**
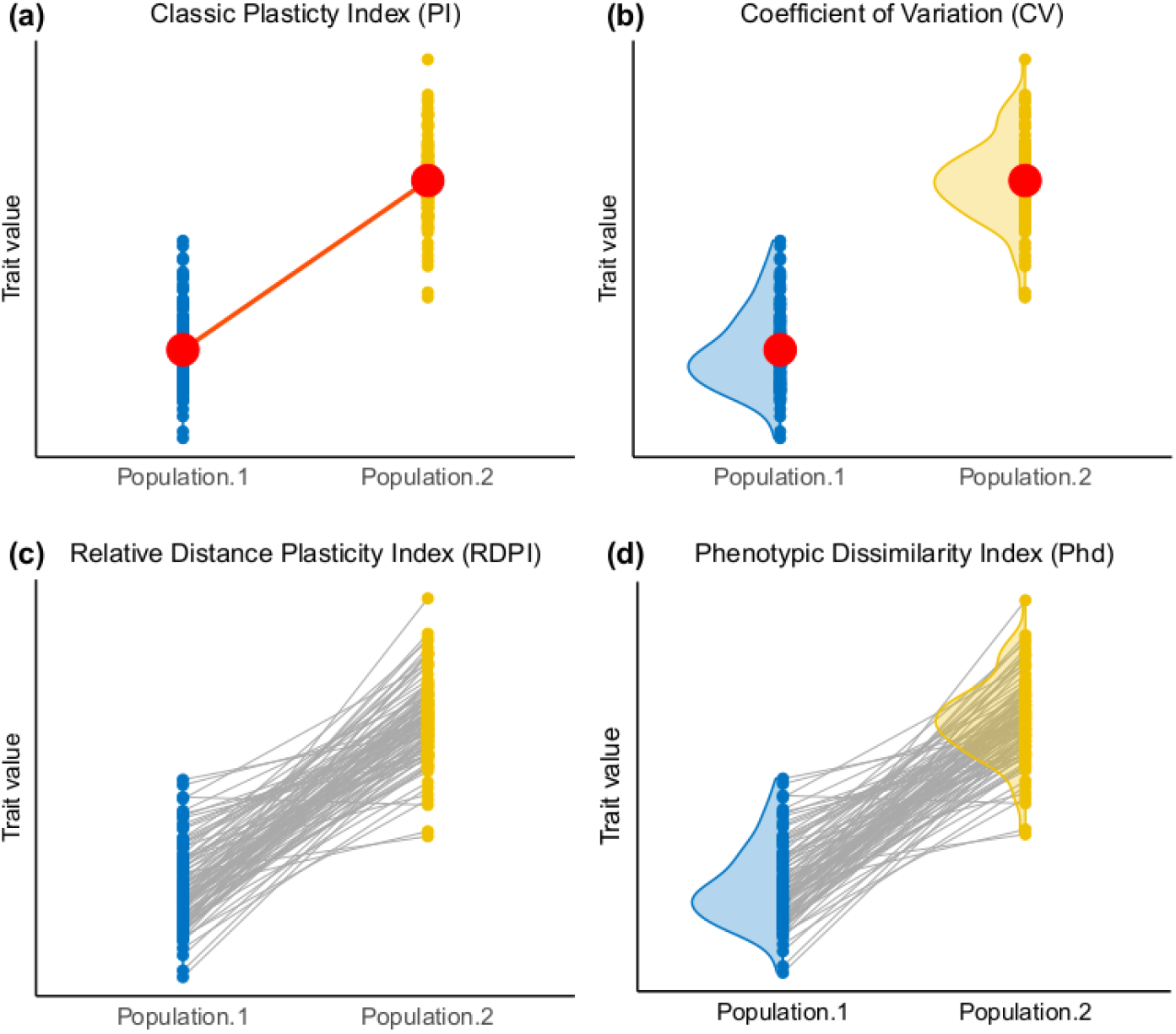
Representation of the calculation of the major estimators of phenotypic variation available in the literature as compared to the Phenotypic Dissimilarity Index (PhD) proposed in this paper. **(a)** Classic Plasticity Index (PI, Valladares et al. 2000) – only trait means within populations are used to calculate PI (red dots) between populations. **(b)** Coefficient of Variation (CV) – mean and within population trait variability (shaded areas) of a trait are used for CV calculation within each considered population, but CV cannot be partitioned between populations. **(c)** Relative Distance Plasticity Index (RDPI) – it calculates the mean phenotypic dissimilarity among all the individuals from different populations (grey lines) without taking into account within population trait variability. **(d)** PhD index – it calculates the mean phenotypic dissimilarity among all the individuals from different populations (grey lines) by taking into account within population trait variability (shaded areas).

By building on the RDPI proposed by Valladares et al. (2006), we aimed to formulate a *PVar* that we call ‘Phenotypic Dissimilarity Index’ (PhD), able to quantify ITV between populations or groups of individuals growing in different environmental conditions by accounting for the phenotypic variability within each population/group (**Fig. 1d**).

PhD can also be used to quantify ITV across individuals within a single site while accounting for intraindividual variability. PhD, as the other *PVar* indices, is bounded between 0 and 1, making its interpretation and comparison across species/individuals straightforward. As such, the proposed PhD index represents an updated framework to estimate phenotypic variation by simultaneously accounting for different ITV components depending on the comparison being made.

We provide two worked examples to show that:

1. Accounting for within population variability is essential when calculating *PVar* between populations. The same applies when addressing interindividual variability within a site while controlling for within-individual trait variance;
2. PhD provides reliable measures of *PVar* – i.e. PhD value increases with environmental dissimilarity among sites (**Supporting Information Fig. S1**) – and it is qualitatively comparable with previous *PVar* indexes. However, since previous *PVar* indexes do not account for intrapopulation/intragroup variability, they consistently return less parsimonious *PVar* estimates compared to the proposed PhD index.

We propose the PhD index as a good candidate to effectively estimate *PVar* between populations/groups, and to compare *PVar* among species. The R function to calculate the PhD index is provided in **Appendix S1**.

## Methods

### Index formulation

Studies focused on the phenotypic variation (*PVar*) within a species are usually based on the evaluation of how a target trait changes across a set of varying environmental conditions, either experimentally or in the field. For a species that occupies *K* positions along an environmental gradient or levels of a treatment (henceforth, such positions or treatments are referred to as environmental states), let *N*_*k*_ be the number of individuals in environmental state *k* (*k* = 1, 2,…, *K*) and *τ*_*ik*_ be the value of trait *τ* for individual *i* (*i* = 1, 2,…, *N*_*k*_) in the environmental state *k*.

According to Valladares et al. (2006), in the simplest case of only two environmental states, *k* and *m*, we can summarize the phenotypic variation of trait *τ* as the expected trait dissimilarity between two individuals drawn at random, one from each environmental state:

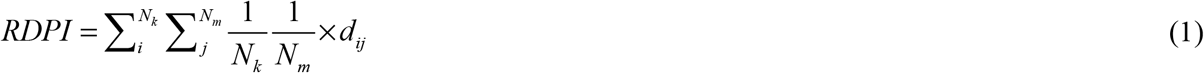

where 1 /*N*_*k*_ is the probability of drawing individual *i* from state *k*, 1 /*N*_*m*_ is the probability of drawing individual *j* from state *m* and *d*_*ij*_ is any symmetric dissimilarity measure between the trait values of individuals *i* and *j* such that *d*_*ij*_ = *d*_*ji*_ and *d*_*ii*_ = 0. For one single trait *τ*, Valladares et al. (2006) proposed to calculate *d*_*ij*_ as *d*_*ij*_ = |*τ*_*ik*_ −*τ*_*jk*_ |*/*(*τ*_*ik*_ +*τ*_*jk*_). This measure is basically the univariate version of the Bray & Curtis (1957) dissimilarity, a multivariate dissimilarity index that has been extensively used by ecologists. With this index, since *d*_*ij*_ is bounded between zero and one, RDPI is also bounded in the same range.

Note that for sake of generality, in Eq. (1) we assume that the number of individuals in environmental states *k* and *m* does not necessarily has to be the same. Note also that by replacing the univariate measure *d*_*ij*_ with one of the many available multivariate dissimilarity measures in the ecologist toolbox, we can easily generalize the calculation of *PVar* to multiple traits of various statistical types - e.g. nominal, fuzzy, ordinal (Pavoine et al., 2009; Legendre and Legendre 2012).

In the context of biodiversity theory, the same index was independently proposed by Rao (1982) and by Webb et al. (2008) to measure the functional or phylogenetic dissimilarity between two species assemblages. However, Rao (1982) noted that Eq. (1) (i.e. RDPI) cannot be immediately used as a measure of phenotypic variation between environmental states *k* and *m*. This is because the value of RDPI depends on the trait variability within each environmental state, which can be defined as:

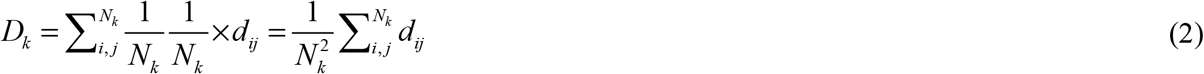

*D*_*k*_ thus summarizes the expected trait dissimilarity between two individuals *i* and *j* drawn at random from environmental state *k*.

Here, it is worth noting that if the trait dissimilarity *d*_*ij*_ among the individuals in *k* is calculated as half the squared Euclidean distance of trait *τ* between individuals *i* and *j* □ i.e. if 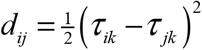- then *D*_*k*_ is equal to the variance of *τ* for all individuals in *k* (Pavoine, 2012). Accordingly, with the Rao-Valladares framework, within-environmental state phenotypic variation *D*_*k*_ is directly related to the variance of the target trait *τ*.

The consequence of the dependence of RDPI from *D*_*k*_ and *D*_*m*_ is that RDPI violates the basic condition that for two identical environmental states with the same number of individuals in each state and identical trait distribution - i.e. if for each individual *i* in environmental state *k* there is an individual *j* in *m* such that *τ*_*ik*_ = *τ*_*jm*_ -the measure take the value zero. In other words, if we compare a given environmental state with itself, the resulting value of RDPI can be larger than zero, thus violating the intuitive assumption that a measure of trait dissimilarity for two identical environmental states cannot be larger than zero (Pavoine & Ricotta, 2014). In this instance, it is in fact possible to have different values of RDPI depending on the distribution of trait values within each environmental state.

To overcome this problem, Rao (1982) demonstrated that if the dissimilarity matrix **D** with elements *d*_*ij*_ is squared Euclidean we have 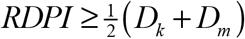. A dissimilarity matrix **D** with elements *d*_*ij*_ is said to be squared Euclidean, if the associated dissimilarity matrix **Δ** with elements 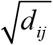 is Euclidean so that **Δ** can be associated with clouds of points in Euclidean space without distortions (Gower & Legendre, 1986). Accordingly, for a squared Euclidean dissimilarity coefficient *d*_*ij*_ bounded in the range [0, 1], Pavoine & Ricotta (2014) proposed the following normalized version of the RDPI index:

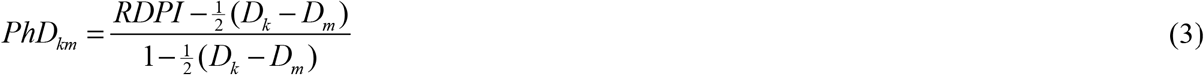

where the acronym PhD stands for Phenotypic Dissimilarity, an estimator of *PVar* that expresses the expected trait dissimilarity across environmental states in the range [0, 1] independently of the trait variability within each environmental state.

In Eq. (3), *PhD*_*km*_ is obtained by linearly rescaling RDPI between its minimum and maximum value *PhD*_*km*_ = (*RDPI* − min _*RDPI*_) / (max_*RDPI*_ − min_*RDPI*_). As such, PhD and RDPI converge in their *PVar* estimates only when the variability of a trait within each environmental state is zero (**Fig. 2a**), and increasingly diverge as the trait standard deviation within environmental states increases (**Fig. 2b-d**).

**Fig. 2.**
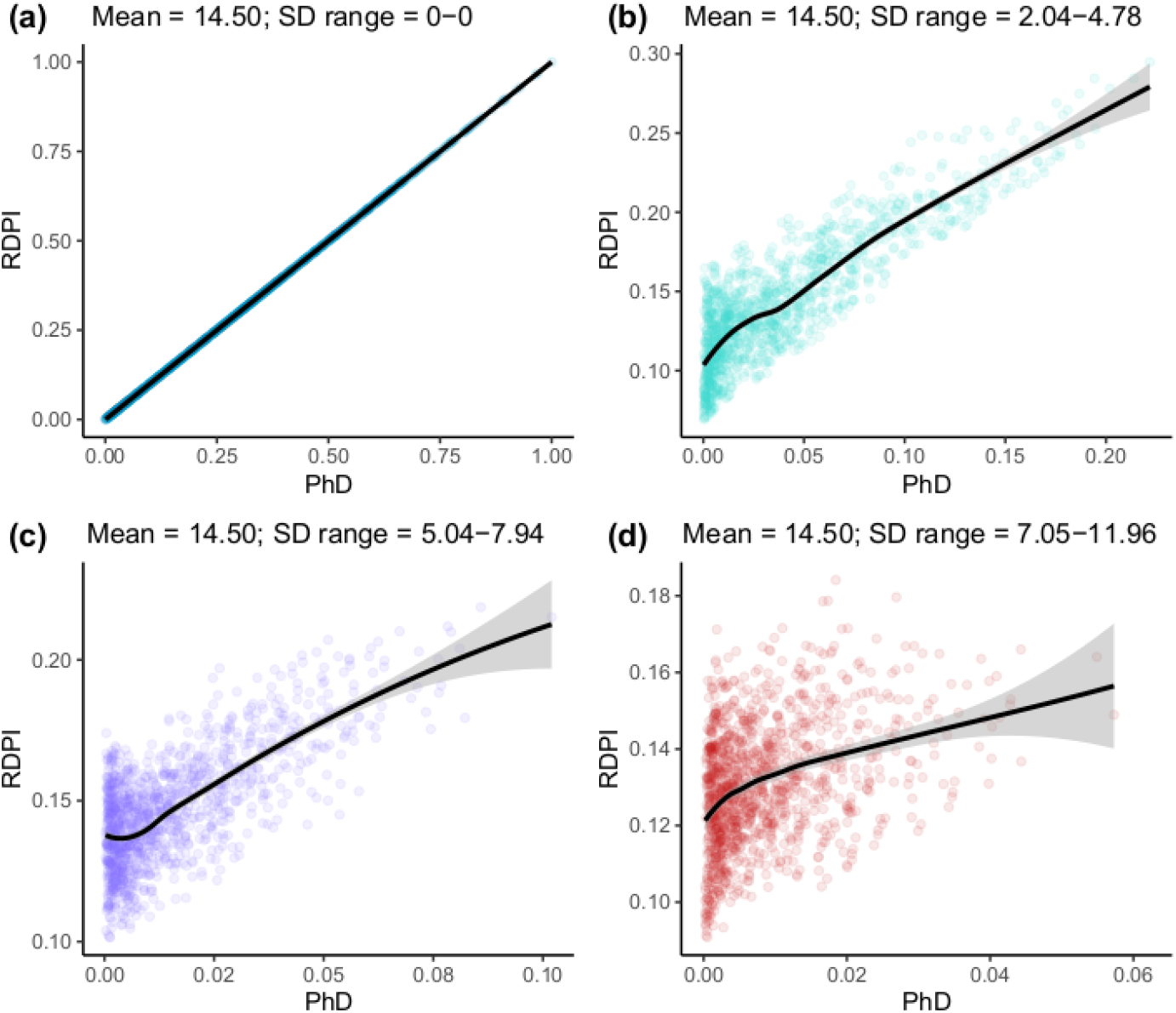
Relationship between Relative Distance Plasticity Index (RDPI) and the Phenotypic Dissimilarity Index (PhD). In each panel, the relationship was evaluated among simulated populations (each data point) having exactly the same mean value (14.50) of a trait but varying standard deviation in the range **(a)** 0-0, **(b)** 2.04-4.78, **(c)** 5.04-7.94 and **(d)** 7.05-11.96. The black solid line is a Loess fit. The shaded area represents the confidence intervals around the Loess fit.

Note here that the trait dissimilarity between pairs of individuals originally proposed by Valladares et al. (2006) *d*_*ij*_ = | *τ*_*ik*_ − *τ*_*jm*_ | */*(*τ*_*ik*_ + *τ*_*jm*_) is squared Euclidean (Legendre and Legendre 2012). Therefore, it can be used to calculate *PhD*_*km*_ in a meaningful way.

Finally, a simple and intuitive way to generalize *PhD*_*km*_ to more than two environmental states, which is usually adopted in community ecology for calculating the beta diversity of a set of species assemblages, consists in calculating the mean value of *PhD*_*km*_ for all possible *K* (*K* −1)/2 pairs of environmental states (e.g. Legendre & De Cáceres, 2013):

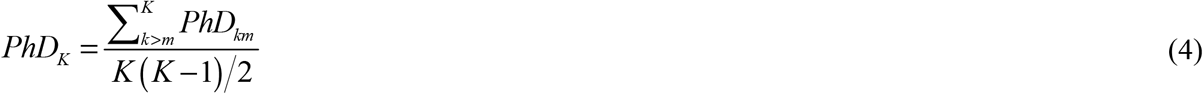

PhD_K_ thus represents a generalization of Eq. (3) to any number of environmental states.

### Example 1: Testing the effect of interindividual variability in *PVar* estimates using simulated data

By controlling for interindividual variability when comparing populations/groups across environmental states, the PhD index is independent on the trait variance within environmental states. We use an example with simulated data to display such desirable property of the PhD index. As already described, the PhD index has two integrated components: the first is a dissimilarity estimator among individuals belonging to different environmental states, which corresponds with the RDPI component in Eq. 3 (*Between* component from now on); the second, which summarizes trait variability within each considered environmental state, corresponds with the D term in Eq. 3 (*Within* component). Because we want to display the behavior of PhD index (*Between* - *Within*) as compared to its *Between* component alone, we generated four scenarios (**Fig. 3a-d**) each including 4 populations (i.e. 6 contrasts), all of them with the same mean trait value but with trait variance changing across levels, as follows:

**Fig. 3.**
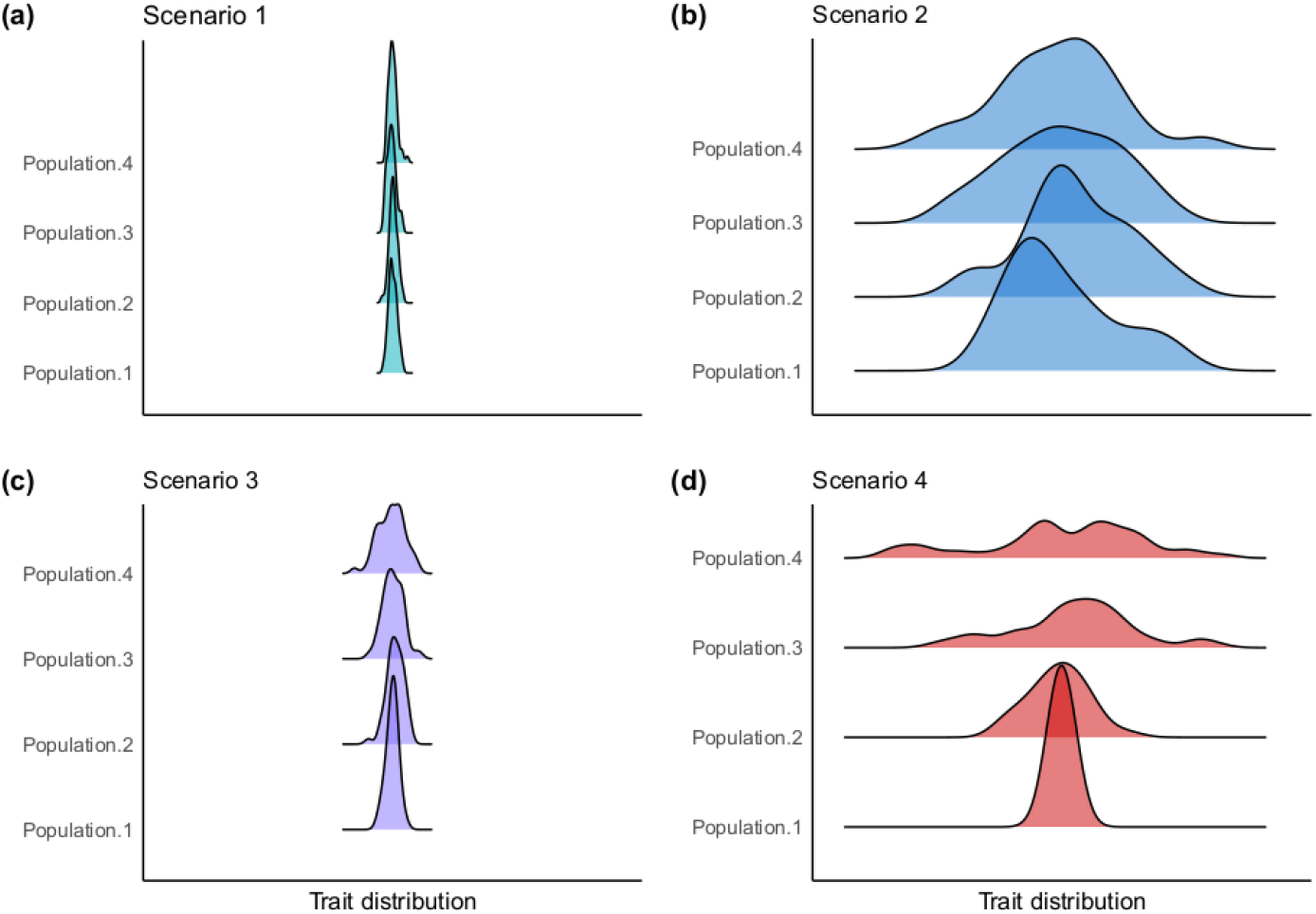
The four simulated scenarios used in Example 1. In each scenario all the populations have the same mean but in **(a)** Scenario 1 - the trait variance is homogeneous across populations and low in magnitude within each population. **(b)** Scenario 2 - the trait variance is homogeneous across populations and high in magnitude within each population. **(c)** Scenario 3 - the trait variance is heterogeneous across populations but relatively low in magnitude within each population. **(d)** Scenario 4 - the trait variance is heterogeneous across populations but relatively high in magnitude within each population.

1. ***Scenario 1***: within-population trait variance is low and of similar magnitude across all populations (**Fig. 3a**). We expected the PhD index and the *Between* component to behave similarly given the low magnitude and similar value of the virtual trait variance within each population. In other words, the *D* component of the PhD index should be negligible, rendering the PhD similar to the RDPI. In this scenario, the two indexes should be highly correlated.
2. ***Scenario 2***: within-population trait variance is high and of similar magnitude across all populations (**Fig. 3b**). As the only difference with *Scenario 1* is just an increase in the magnitude of within-population trait variance, we expected the PhD index and the RDPI to be still correlated.
3. ***Scenario 3***: within-population trait variance is relatively low across all populations, but not all populations have the same trait variance (**Fig. 3c**). Under this scenario, we expected the indexes to start losing their correlation, as the differences in the virtual trait distribution within populations should decouple PhD index estimates from that of the RDPI (see Index formulation section and **Fig 2a-d**).
4. ***Scenario 4***: within-population trait variance is relatively high across all populations, but not all populations have the same trait variance (**Fig. 3d**). Similar to Scenario 3, we expected the two indexes to completely lose their correlation.

### Example 2: Comparing PhD index with other *PVar* estimators

For this example, we used an already published dataset of some Mediterranean grassland species growing along a topographical and soil water content gradient (Carmona et al., 2015). From this dataset, we used specific leaf area (SLA) measurements. Following Valladares et al. (2006), we selected the following *PVar*: i) slope of the linear regression of a trait *vs*. soil water content (*Slope*) (Schlichting & Pigliucci, 1998); ii) coefficient of variation (CV); iii) classic plasticity index (PI); and iv) Relative Distance Plasticity Index (RDPI) (see Introduction and **Fig. 1a-c** for details on PI, CV and RDPI calculations). For this example, we selected only species that were present in at least four plots along the considered gradient, as we considered four plots to be the minimum number data points allowed to calculate a *Slope* estimator via simple linear regression analysis (e.g. Veresoglou & Peñuelas, 2019). Simple linear regression analysis refers to the relationship SLA *vs*. Soil Water Content. Seventeen species were included in the final analysis, and we used such species to calculate the PhD index and the other four *PVar* across plots representing species position along the soil water content gradient. The final dataset included species-specific mean values of PhD, Slope, CV, PI and RDPI calculated across the gradient. We explored the relationship between PhD index and the other four *PVar* estimators using simple linear regression analysis.

## Results

### Example 1: Testing the effect of interindividual variability in *PVar* estimates using simulated data

The results of Example 1 show that when within-population variance is either high or heterogeneous across populations, the results provided by the PhD index and by RDPI rapidly diverge. In particular, we found that the two indexes return comparable *PVar* estimates (R = 0.90; p □ 0.05) only when the trait variance within each population is low in magnitude and similar across populations (Scenario 1, **Fig. 3a**). However, when the variance differed both within and between populations (Scenarios 3 and 4, **Fig. 3c-d**), the two indexes were uncorrelated with R ranging between −0.04 and 0.16 in Scenario 3 and 4 (p always □ 0.05), independently of the variance magnitude. Surprisingly, when increasing the magnitude of the within population variance while keeping the variance magnitude comparable across populations - i.e. Scenario 2 (**Fig. 3b**) - the indexes turned out to be negatively correlated (R = −0.65; p □ 0.05), even if we expected Scenario 2 to yield similar results as Scenario 1. This can be explained considering that the four simulated populations in Scenario 2 slightly differed among them in terms of trait variance, but such very small differences, arising due to the simulation procedure, are enough to let the indexes to diverge when the trait variance within population is high.

### Example 2: Comparing PhD index with other *PVar* estimators

Results of this analysis (**Fig. 4a-d**) show that PhD index is correlated with three of the four considered *PVar* estimators. The only exception is the relationship PhD-*Slope* that was not significant because trait variability along the gradient is not fully linear (**Fig. S2**; Carmona et al., 2015), and *Slope*-based *PVar* estimates might not be reliable. At any rate, PhD was correlated with *CV, PI* and *RDPI* (**Fig. 4b-d**), demonstrating that PhD estimates are qualitatively comparable with those from other estimators. However, the estimates provided by the PhD index are generally smaller than that of *CV, PI* and *RDPI*, as all the data points in **Fig. 4b-d** fall above the 1:1 line.

**Fig. 4.**
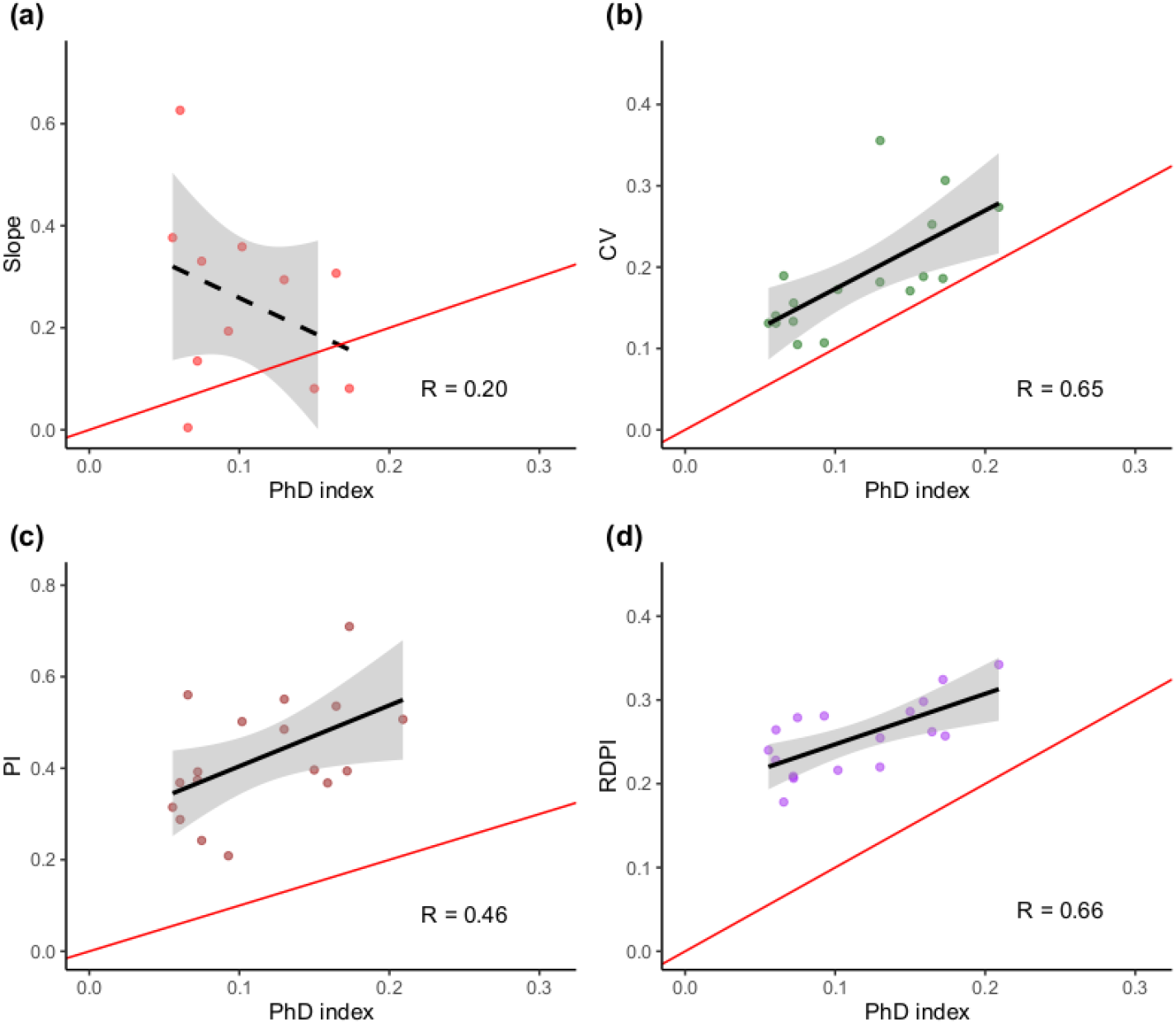
Relationship among Phenotypic Dissimilarity Index (PhD) and common estimators of phenotypic plasticity. **(a)** Slope of reaction norm (Slope); **(b)** coefficient of variation (CV); **(c)** classic Plasticity Index (PI, *sensu* Valladares et al., 2000); **(d)** Relative Distance Plasticity Index (RDPI, Valladares et al., 2006). See main text for further details on the selected phenotypic plasticity estimators. Solid line indicate significant relationship at p < 0.05. Red line represents a 1:1 relationship. Correlation coefficients (R) are also shown.

## Discussion

The proposed PhD index quantifies phenotypic variability, or ITV, or phenptypic plasticity (*sensu* Valladares et al., 2000) across populations of a species by accounting for the effect of interindividual trait variability. The results from our simulation example demonstrate the importance of using the PhD index when one wants to quantify intraspecific variability between-populations using *PVar* estimators where the within-population trait variability cannot be considered negligible. Within-population trait variability can be considered negligible when it tends to zero, but this is very rare in most ecological studies. Moreover, we have showed that the estimates provided by the PhD index are generally smaller than that of the most common *PVar* estimators (**Fig. 4a-d**). This aspect is of particular interest as *PVar* estimates are classically used to identify traits that are more responsive to environmental gradients, and these traits are usually interpreted as drivers of species responses to environmental changes, including global climate change. Our results show that such conclusions need to be re-analyzed in view of the within-population variability effect on *PVar* across environmental gradients, and the PhD index represents an essential tool to reach such an overarching objective.

The PhD index we propose is insensitive to the within-population interindividual trait variability when the target is quantifying phenotypic variability between populations. When does accounting for within-population interindividual trait variability matters? Interindividual variability within populations is a central topic in ecology and evolution, because of its relevance in determining, for instance, the relaxation of intra- and interspecific competition through habitat specialization (e.g. Araújo et al., 2011; Bolnick et al., 2010; Cam et al., 2002; Devictor et al., 2010; Roughgarden, 1972; Violle et al., 2012). Each individual in a population can in fact use a subset of the available resources in a ‘population’s niche’ (Bolnick et al., 2003), and this can shape individuals’ resource acquisition and use strategies accordingly. Individual specialization within-populations increases the variance of traits linked to the acquisition and the use of alternative resources, and such trait variance can ultimately differ between populations arranged along an environmental gradient. Intra- and inter-population niche variation is in fact considered as an important target for natural selection (Bolnick et al., 2003) as single individuals of the same species can be always subjected to contrasting selective pressures. Therefore, we argue that the proposed PhD index is an essential tool to quantify inter-population phenotypic variability in the field, where within-population trait variance is surely relevant. Besides, by returning measures of intraspecific variability bounded between 0 and 1, and standardized for within-population trait variability, the PhD index can be used to easily compare phenotypic variability between species across an environmental gradient. The PhD index also provides an alternative to classic analyses such as ANOVA or PERMANOVA, whose applicability is constrained by the response variable distribution/dispersion, and their results, expressed in variance units, cannot be always intuitively compared across species.

Common practice in functional ecology is to use a set of standardized rules for sampling individuals’ features in order to obtain estimates of functional traits (e.g. Pérez-Harguindeguy et al., 2016 for plants). Despite such standardization of sampling procedures, for the reasons outlined above, traits sampled within a species have an inherent variability (Albert et al., 2011). Our index indeed allows the user to account for trait variability at different levels, permitting to summarize between-populations differences in a consistent and comparable way across species as the determination of *PVar* will not anymore depend on the interindividual variability within each of the considered populations. This also applies when quantifying *PVar* within populations by accounting for intraindividual variability.

In sum, we showed that the PhD index is a phenotypic variability estimator that reaches the greatest level of generality compared to other estimators. In particular, it represents a generalization of the RDPI of Valladares et al. (2006). We also argue that the PhD index properties will become especially relevant when trying to determine the key traits underlying a species response to contrasting environmental conditions, particularly in a climate change context.

## Supporting information

Appendixes

## Acknowledgements

This study was financed by the Estonian University of Life Sciences (grant number: P200187PKEL and P200190PKEL, for GP and LL respectively), the Estonian Ministry of Education and Research (PSG293 for CPC), and the Sapienza University of Rome (RM11916B6A2EA7D5 and RM120172AF29E651 for CR and LV, respectively).

## Authors’ contributions statement

GP and CR conceived the idea; CR formulated the index; CPC programmed the R script; GP and CPC planned the data analysis; GP lead the data analysis; LL and LV contributed to conceptualization, literature review and results interpretation. GP wrote the first draft of the manuscript. All authors contributed critically to the drafts and gave final approval for publication.

## References

Albert, C. H., Grassein, F., Schurr, F. M., Vieilledent, G., & Violle, C. (2011). When and how should intraspecific variability be considered in trait-based plant ecology? Perspectives in Plant Ecology, Evolution and Systematics, 13(3), 217–225. https://doi.org/10.1016/j.ppees.2011.04.003

Araújo, M. S., Bolnick, D. I., & Layman, C. A. (2011). The ecological causes of individual specialisation. Ecology Letters, 14(9), 948–958. https://doi.org/10.1111/j.1461-0248.2011.01662.x

Arnold, P. A., Kruuk, L. E. B., & Nicotra, A. B. (2019). How to analyse plant phenotypic plasticity in response to a changing climate. New Phytologist, 222(3), 1235–1241. https://doi.org/10.1111/nph.15656

Balaguer, L., Martínez-Ferri, E., Valladares, F., Pérez-Corona, M. E., Baquedano, F. J., Castillo, F. J., & Manrique, E. (2001). Population Divergence in the Plasticity of the Response of Quercus coccifera to the Light Environment. Functional Ecology, 15(1), 124–135.

Bolnick, D. I., Ingram, T., Stutz, W. E., Snowberg, L. K., Lau, O. L., & Paull, J. S. (2010). Ecological release from interspecific competition leads to decoupled changes in population and individual niche width. Proceedings of the Royal Society B: Biological Sciences, 277(1689), 1789–1797. https://doi.org/10.1098/rspb.2010.0018

Bolnick, D. I., Svanbäck, R., Fordyce, J. A., Yang, L. H., Davis, J. M., Hulsey, C. D., & Forister, M. L. (2003). The Ecology of Individuals: Incidence and Implications of Individual Specialization. The American Naturalist, 161(1), 1–28. https://doi.org/10.1086/343878

Bray, J. R., & Curtis, J. T. (1957). An Ordination of the Upland Forest Communities of Southern Wisconsin. Ecological Monographs, 27(4), 325–349. https://doi.org/10.2307/1942268

Cam, E., Link, W. A., Cooch, E. G., Monnat, J., & Danchin, E. (2002). Individual Covariation in Life□History Traits: Seeing the Trees Despite the Forest. The American Naturalist, 159(1), 96– 105. https://doi.org/10.1086/324126

Carmona, C. P., Rota, C., Azcárate, F. M., & Peco, B. (2015). More for less: Sampling strategies of plant functional traits across local environmental gradients. Functional Ecology, 29(4), 579– 588. https://doi.org/10.1111/1365-2435.12366

Castro-Díez, P., Navarro, J., Pintado, A., Sancho, L. G., & Maestro, M. (2006). Interactive effects of shade and irrigation on the performance of seedlings of three Mediterranean Quercus species. Tree Physiology, 26(3), 389–400. https://doi.org/10.1093/treephys/26.3.389

de Bello, F., Lavorel, S., Albert, C. H., Thuiller, W., Grigulis, K., Dolezal, J., Janeček, Š., & Lepš, J. (2011). Quantifying the relevance of intraspecific trait variability for functional diversity. Methods in Ecology and Evolution, 2(2), 163–174. https://doi.org/10.1111/j.2041-210X.2010.00071.x

Devictor, V., Clavel, J., Julliard, R., Lavergne, S., Mouillot, D., Thuiller, W., Venail, P., Villéger, S., & Mouquet, N. (2010). Defining and measuring ecological specialization. Journal of Applied Ecology, 47(1), 15–25. https://doi.org/10.1111/j.1365-2664.2009.01744.x

Garnier, E., Laurent, G., Bellmann, A., Debain, S., Berthelier, P., Ducout, B., Roumet, C., & Navas, M.-L. (2001). Consistency of species ranking based on functional leaf traits. New Phytologist, 152(1), 69–83. https://doi.org/10.1046/j.0028-646x.2001.00239.x

Garnier, E., Navas, M.-L., & Grigulis, K. (2015). Plant Functional Diversity: Organism traits, community structure, and ecosystem properties. Oxford University Press. https://doi.org/10.1093/acprof:oso/9780198757368.001.0001

Gower, J. C., & Legendre, P. (1986). Metric and Euclidean properties of dissimilarity coefficients. Journal of Classification, 3(1), 5–48. https://doi.org/10.1007/BF01896809

Henn, J. J., Buzzard, V., Enquist, B. J., Halbritter, A. H., Klanderud, K., Maitner, B. S., Michaletz, S. T., Pötsch, C., Seltzer, L., Telford, R. J., Yang, Y., Zhang, L., & Vandvik, V. (2018). Intraspecific Trait Variation and Phenotypic Plasticity Mediate Alpine Plant Species Response to Climate Change. Frontiers in Plant Science, 9. https://doi.org/10.3389/fpls.2018.01548

Herrera, C. M., Medrano, M., & Bazaga, P. (2015). Continuous within-plant variation as a source of intraspecific functional diversity: Patterns, magnitude, and genetic correlates of leaf variability in Helleborus foetidus (Ranunculaceae). American Journal of Botany, 102(2), 225–232. https://doi.org/10.3732/ajb.1400437

Kuppler, J., Albert, C. H., Ames, G. M., Armbruster, W. S., Boenisch, G., Boucher, F. C., Campbell, D. R., Carneiro, L. T., Chacón-Madrigal, E., Enquist, B. J., Fonseca, C. R., Gómez, J. M., Guisan, A., Higuchi, P., Karger, D. N., Kattge, J., Kleyer, M., Kraft, N. J. B., Larue-Kontić, A.- A. C., … Junker, R. R. (2020). Global gradients in intraspecific variation in vegetative and floral traits are partially associated with climate and species richness. Global Ecology and Biogeography, 29(6), 992–1007. https://doi.org/10.1111/geb.13077

Legendre, P., & De Cáceres, M. (2013). Beta diversity as the variance of community data: Dissimilarity coefficients and partitioning. Ecology Letters, 16(8), 951–963. https://doi.org/10.1111/ele.12141

Legendre, P., Legendre, L. (2012) Numerical Ecology. Elsevier, Amsterdam.

Mason, N. W. H., Orwin, K. H., Lambie, S., Waugh, D., Pronger, J., Carmona, C. P., & Mudge, P. (2020). Resource-use efficiency drives overyielding via enhanced complementarity. Oecologia, 193(4), 995–1010. https://doi.org/10.1007/s00442-020-04732-7

McGill, B. J., Enquist, B. J., Weiher, E., & Westoby, M. (2006). Rebuilding community ecology from functional traits. Trends in Ecology & Evolution, 21(4), 178–185. https://doi.org/10.1016/j.tree.2006.02.002

Nicotra, A. B., Atkin, O. K., Bonser, S. P., Davidson, A. M., Finnegan, E. J., Mathesius, U., Poot, P., Purugganan, M. D., Richards, C. L., Valladares, F., & van Kleunen, M. (2010). Plant phenotypic plasticity in a changing climate. Trends in Plant Science, 15(12), 684–692. https://doi.org/10.1016/j.tplants.2010.09.008

Niinemets, Ü., Keenan, T. F., & Hallik, L. (2015). A worldwide analysis of within-canopy variations in leaf structural, chemical and physiological traits across plant functional types. New Phytologist, 205(3), 973–993. https://doi.org/10.1111/nph.13096

Niu, K., Zhang, S., & Lechowicz, M. J. (2020). Harsh environmental regimes increase the functional significance of intraspecific variation in plant communities. Functional Ecology, 34(8), 1666– 1677. https://doi.org/10.1111/1365-2435.13582

Pavoine, S. (2012). Clarifying and developing analyses of biodiversity: Towards a generalisation of current approaches. Methods in Ecology and Evolution, 3(3), 509–518. https://doi.org/10.1111/j.2041-210X.2011.00181.x

Pavoine, S., & Ricotta, C. (2014). Functional and phylogenetic similarity among communities. Methods in Ecology and Evolution, 5(7), 666–675. https://doi.org/10.1111/2041-210X.12193

Pavoine, S., Vallet, J., Dufour, A.-B., Gachet, S., & Daniel, H. (2009). On the challenge of treating various types of variables: Application for improving the measurement of functional diversity. Oikos, 118(3), 391–402. https://doi.org/10.1111/j.1600-0706.2008.16668.x

Pérez-Harguindeguy, N., Díaz, S., Garnier, E., Lavorel, S., Poorter, H., Jaureguiberry, P., Bret-Harte, M. S., Cornwell, W. K., Craine, J. M., Gurvich, D. E., Urcelay, C., Veneklaas, E. J., Reich, P. B., Poorter, L., Wright, I. J., Ray, P., Enrico, L., Pausas, J. G., Vos, A. C. de, … Cornelissen, J. H. C. (2016). Corrigendum to: New handbook for standardised measurement of plant functional traits worldwide. Australian Journal of Botany, 64(8), 715–716. https://doi.org/10.1071/bt12225_co

Poorter, H., Jagodzinski, A. M., Ruiz-Peinado, R., Kuyah, S., Luo, Y., Oleksyn, J., Usoltsev, V. A., Buckley, T. N., Reich, P. B., & Sack, L. (2015). How does biomass distribution change with size and differ among species? An analysis for 1200 plant species from five continents. New Phytologist, 208(3), 736–749. https://doi.org/10.1111/nph.13571

Puglielli, G. (2019). Beyond the Concept of Winter-Summer Leaves of Mediterranean Seasonal Dimorphic Species. Frontiers in Plant Science, 10, 696. https://doi.org/10.3389/fpls.2019.00696

Puglielli, G., Catoni, R., Spoletini, A., Varone, L., & Gratani, L. (2017). Short-term physiological plasticity: Trade-off between drought and recovery responses in three Mediterranean Cistus species. Ecology and Evolution, 7(24), 10880–10889. https://doi.org/10.1002/ece3.3484

Puglielli, G., Laanisto, L., Poorter, H., & Niinemets, Ü. (2021). Global patterns of biomass allocation in woody species with different tolerances of shade and drought: Evidence for multiple strategies. The New Phytologist, 229(1), 308–322. https://doi.org/10.1111/nph.16879

Puglielli, G., Varone, L., & Gratani, L. (2019). Diachronic adjustments of functional traits scaling relationships to track environmental changes: Revisiting Cistus species leaf cohort classification. Flora, 254, 173–180. https://doi.org/10.1016/j.flora.2018.08.010

Rao, C. R. (1982). Diversity and dissimilarity coefficients: A unified approach. Theoretical Population Biology, 21(1), 24–43. https://doi.org/10.1016/0040-5809(82)90004-1

Roughgarden, J. (1972). Evolution of Niche Width. The American Naturalist, 106(952), 683–718. https://doi.org/10.1086/282807

Rutherford, S., Bonser, S. P., Wilson, P. G., & Rossetto, M. (2017). Seedling response to environmental variability: The relationship between phenotypic plasticity and evolutionary history in closely related Eucalyptus species. American Journal of Botany, 104(6), 840–857. https://doi.org/10.3732/ajb.1600439

Sandquist, D. R., & Ehleringer, J. R. (1997). Intraspecific variation of leaf pubescence and drought response in Encelia farinosa associated with contrasting desert environments. The New Phytologist, 135(4), 635–644. https://doi.org/10.1046/j.1469-8137.1997.00697.x

Schlichting, C., & Pigliucci, M. (1998). Phenotypic Evolution: A Reaction Norm Perspective. Sinauer.

Siefert, A., Violle, C., Chalmandrier, L., Albert, C. H., Taudiere, A., Fajardo, A., Aarssen, L. W., Baraloto, C., Carlucci, M. B., Cianciaruso, M. V., de L Dantas, V., de Bello, F., Duarte, L. D. S., Fonseca, C. R., Freschet, G. T., Gaucherand, S., Gross, N., Hikosaka, K., Jackson, B., … Wardle, D. A. (2015). A global meta-analysis of the relative extent of intraspecific trait variation in plant communities. Ecology Letters, 18(12), 1406–1419. https://doi.org/10.1111/ele.12508

Turner, N. C., Schulze, E.-D., Nicolle, D., Schumacher, J., & Kuhlmann, I. (2008). Annual rainfall does not directly determine the carbon isotope ratio of leaves of Eucalyptus species. Physiologia Plantarum, 132(4), 440–445. https://doi.org/10.1111/j.1399-3054.2007.01027.x

Valladares, F., Sanchez-Gomez, D., & Zavala, M. A. (2006). Quantitative estimation of phenotypic plasticity: Bridging the gap between the evolutionary concept and its ecological applications. Journal of Ecology, 94(6), 1103–1116. https://doi.org/10.1111/j.1365-2745.2006.01176.x

Valladares, F., Wright, S. J., Lasso, E., Kitajima, K., & Pearcy, R. W. (2000). Plastic Phenotypic Response to Light of 16 Congeneric Shrubs from a Panamanian Rainforest. Ecology, 81(7), 1925–1936. https://doi.org/10.1890/0012-9658(2000)081[1925:PPRTLO]2.0.CO;2

Veresoglou, S. D., & Peñuelas, J. (2019). Variance in biomass-allocation fractions is explained by distribution in European trees. New Phytologist, 222(3), 1352–1363. https://doi.org/10.1111/nph.15686

Violle, C., Enquist, B. J., McGill, B. J., Jiang, L., Albert, C. H., Hulshof, C., Jung, V., & Messier, J. (2012). The return of the variance: Intraspecific variability in community ecology. Trends in Ecology & Evolution, 27(4), 244–252. https://doi.org/10.1016/j.tree.2011.11.014

Webb, C. O., Ackerly, D. D., & Kembel, S. W. (2008). Phylocom: Software for the analysis of phylogenetic community structure and trait evolution. Bioinformatics, 24(18), 2098–2100. https://doi.org/10.1093/bioinformatics/btn358

Wong, M. K. L., & Carmona, C. P. (2021). Including intraspecific trait variability to avoid distortion of functional diversity and ecological inference: Lessons from natural assemblages. Methods in Ecology and Evolution, 12(5), 946–957. https://doi.org/10.1111/2041-210X.13568

