## Appendixes for "An updated framework to account for inter-individual variability when quantifying phenotypic variation"

Supporting Information

**Fig. S1.** Relationship between PhD index and the dissimilarity in soil water content for three species across ˃ 20 plots in Carmona et al. (2015). Only three species were included because they were the only retained ones (among the seventeen mentioned in the main text) that spanned almost the entire gradient. R^2^ and species name per each relationship is also shown.
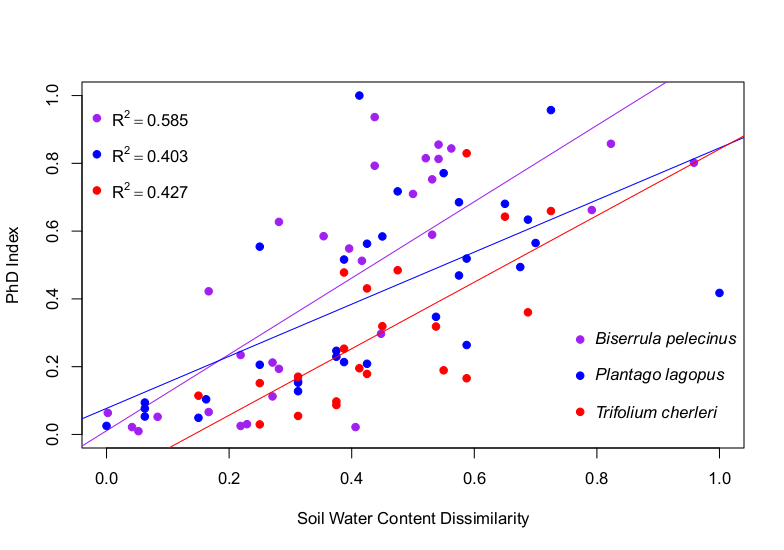

**References**

Carmona, C. P., Rota, C., Azcárate, F. M., & Peco, B. (2015). More for less: Sampling strategies of plant functional traits across local environmental gradients. *Functional Ecology*, *29*(4), 579–588. https://doi.org/10.1111/1365-2435.12366

Appendix S1

*R function to calculate the PhD index*

**Function**

**Description**

PhD computes the PhD index of phenotypic variation, which allows quantifying intraspecific trait variability between populations or groups of individuals growing in different environmental conditions while considering the effect of phenotypic variability within populations or groups.

**Arguments**

***groups***: a vector defining the group (environmental state) to which individuals belongs

***dis***: a matrix of pairwise phenotypic dissimilarity between individuals (bounded between 0 and 1).

**Value**

PhD returns a list containing the following components:

*$between*: dissimilarity estimator among individuals belonging to different environmental states (e.g. populations)

*$PhD*: the value of the PhD index between pairs of environmental states

*$meanPhD*: the mean value of the PhD index across environmental states

```
#Copy and paste the function below in an R script and run it to load it in the global environment
PhD <- function(groups, dis){
  Ngroups <- length(unique(groups))
  raoWithin <- numeric(Ngroups)
  raoBetween <- numeric((Ngroups^2-Ngroups)/2)
  PhD <- numeric((Ngroups^2-Ngroups)/2)
  for(i in 1:Ngroups){
    groupAux<-which(groups == unique(groups)[i])
    disAux <- dis[groupAux, groupAux]
    raoWithin[i]<- sum(disAux)/length(disAux)
    names(raoWithin)[i] <- unique(groups)[i]
  }
  index<-1
  for(i in 1:Ngroups){
    groupAuxi<-which(groups == unique(groups)[i])
    Groupi<-unique(groups)[i]
    for(j in 1:Ngroups){
      if(j>i){
        groupAuxj<-which(groups == unique(groups)[j])
        Groupj<-unique(groups)[j]
        disAux <- dis[groupAuxi, groupAuxj]
        raoBetween[index]<-sum(disAux)/length(disAux)
        names(raoBetween)[index] <- paste0(Groupi, "-", Groupj)

        PhD[index] <- (raoBetween[index] - 0.5*raoWithin[i] - 0.5*raoWithin[j])/
          (1 - 0.5*raoWithin[i] - 0.5*raoWithin[j])
        names(PhD)[index] <- paste0(Groupi, "-", Groupj)
        index<-index+1

      }
    }
  }
  meanPhD <- mean(PhD)
  return(list(PhD = PhD, between = raoBetween,
              meanPhD=meanPhD))
}
#end of function
```

**Example**

*Note: The example below can be replicated only when the function is loaded in the global environment.*

**1)** Defining the number of populations (5 in this case):

nGroups.ex <- 5

**2)** Defining the number of individuals within each population (10 in this case):

nIndGroup.ex <- rep(10, nGroups.ex)

**3)** Naming the 5 populations and the 10 individuals within each population:

```
groups.ex <- numeric()
for(i in 1:nGroups.ex){
  groups.ex<-c(groups.ex, rep(paste0("Population.",i), nIndGroup.ex[i]))
}
names(groups.ex)<- paste0("Ind.", 1:length(groups.ex))
```

**4)** Defining a mean trait value for each population:

meanGroups.ex <- runif(nGroups.ex, 2, 10)

**5)** Defining the standard deviation of the distribution of trait values for each population:

sdGroups.ex <- runif(nGroups.ex, 1, 3)

**6)** Assigning trait values to the individuals using means and standard deviations created at the step 4 and 5, respectively

```
traits.ex <- numeric()
for(i in 1:nGroups.ex){
  traits.ex<-c(traits.ex, rnorm(nIndGroup.ex[i], meanGroups.ex[i], sdGroups.ex[i]))
}
names(traits.ex)<- paste0("Ind.", 1:length(groups.ex))
```

**7)** Calculating the dissimilairty matrix (bound between 0 and 1)

dissim.ex <- as.matrix(dist(traits.ex))
dissim.ex <- dissim.ex / max (dissim.ex) #Bound in the 0-1 range

**8)** Run the PhD function

PhD(groups=groups.ex, dis=dissim.ex)

**Fig. S2.** Mean value of specific leaf area values at each step along the gradient in Carmona et al. (2015).

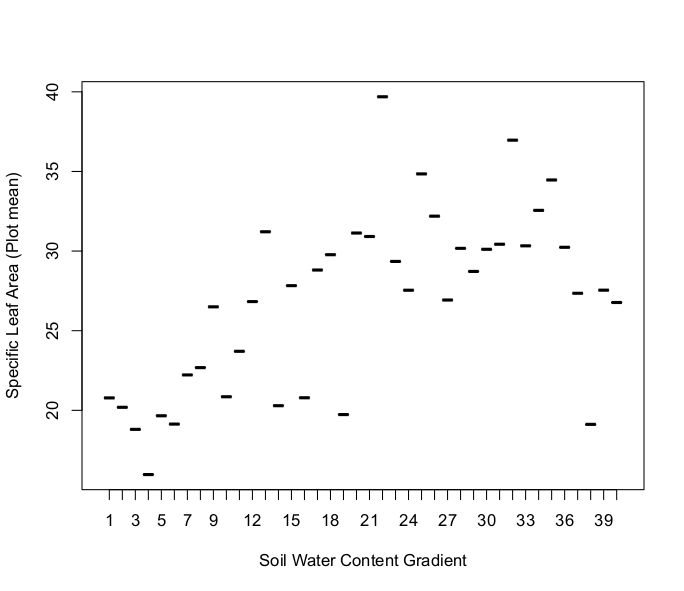

**References**

Carmona, C. P., Rota, C., Azcárate, F. M., & Peco, B. (2015). More for less: Sampling strategies of plant functional traits across local environmental gradients. *Functional Ecology*, *29*(4), 579–588. https://doi.org/10.1111/1365-2435.12366
